# Biophysical, Molecular and Proteomic profiling of Human Retinal Organoids derived Exosomes

**DOI:** 10.1101/2022.04.25.489461

**Authors:** Peggy Arthur, Sangeetha Kandoi, Li Sun, Anil Kalvala, Shallu Kutlehria, Santanu Bhattacharya, Tanmay Kulkarni, Ramesh Nimma, Yan Li, Deepak A. Lamba, Mandip Singh

## Abstract

Extracellular vesicles (EVs) are phospholipid bilayer-bound particles released by cells that play a role in cell-cell communication, signal transduction, and extracellular matrix remodeling. There is a growing interest in EVs for ocular applications as therapeutics, biomarkers, and drug delivery vehicles. EVs secreted from mesenchymal stem cells (MSCs) have shown to provide therapeutic benefits in ocular conditions. However, very little is known about the properties of bioreactors cultured-3D human retinal organoids secreted EVs. This study provides a comprehensive morphological, nanomechanical, molecular, and proteomic characterization of retinal organoid EVs and compares it with human umbilical cord (hUC) MSCs. Nanoparticle tracking analysis indicated the average size of EV as 100–250 nm. Atomic force microscopy showed that retinal organoid EVs are softer and rougher than the hUCMSC EVs. Gene expression analysis by qPCR showed a high expression of exosome biogenesis genes in late retinal organoids derived EVs (>120 days). Immunoblot analysis showed highly expressed exosomal markers Alix, CD63, Flotillin-2, HRS and Hsp70 in late retinal organoids compared to early retinal organoids EVs (<120 days). Protein profiling of retinal organoid EVs displayed a higher differential expression of retinal function-related proteins and EV biogenesis/marker proteins than hUCMSC EVs, implicating that the use of retinal organoid EVs may have a superior therapeutic effect on retinal disorders. This study adds supplementary knowledge on the properties of EVs secreted by retinal organoids and suggests their potential use in the diagnostic and therapeutic treatments for ocular diseases.

## 1. Introduction

Several retinal degenerative diseases, both acquired and inherited, result in permanent vision loss due to the death of the light-sensing photoreceptors in the retina. Retinal neurons, including photoreceptors, are post-mitotic and lack the regenerative potential. Various approaches, including gene therapy, cell transplantation, and optogenetics, are being studied to treat or restore irreversible damage [1–4]. Lack of clinically relevant animal models that closely mimic the human retina’s structure and disease pathogenesis have resulted in the development of *in vitro* retinal models. Human induced pluripotent stem cell (hiPSC) derived retinal organoids serve as an invaluable tool for disease modelling [5, 6], cell therapy [7], drug discovery [8, 9] and for studying human retinogenesis[10–12].

The development of hiPSC technology is a critical milestone in medicine due to its ability to differentiate into 220 different cell types, developing patient-specific disease models and for the treatments for numerous diseases [13–17]. Differentiating photoreceptors using hiPSC holds great potential for better understanding and screening therapeutics for various forms for retinal degenerative diseases including Retinitis Pigmentosa (RP) and age-related macular degeneration (AMD). Novel treatments for retinal diseases associated with the loss of photoreceptors have been developed due to the advent of human embryonic stem cell (hESC) and hiPSC-derived photoreceptors [18].

Extracellular vesicles (EVs), including exosomes, microvesicles, and apoptotic bodies, are lipid bilayer-bound biomolecules secreted by cells, consisting of various nucleic acid and soluble transmembrane proteins [19]. EVs differ in size (30-1000 nm), biogenesis and release pathways, supported functions, and sub-cellular origin [20–22]. Among the EVs, exosomes are spherical, 30-200 nm sized subset, comprising of diverse lipid and protein molecules and are fundamentally secreted by all cell types naturally *via* exocytosis reflecting the physiological state of cells [23]. Although there has been an increasing interest in the therapeutic potential of several cell-secreted EVs for ocular diseases, knowledge in this field is still limited [24]. Multiple studies have explored the biological role of EVs in signal transduction [25], immune regulation [26], repair regeneration [26], biomarkers [27] and as drug delivery vehicles [28].

However, studies to elucidate the biogenesis of EVs from retinal organoids [29] is relatively a new area of interest in exosome theranostics [30]. EVs produced by different cell types or conditions may differ in their content and have varying effects on their target tissues. During disease or injury, retinal astroglial cells-derived EVs have driven choroidal neovascularization, whereas retinal pigment epithelial cells-derived EVs have conferred pro-angiogenic effects when given intravenously and *via* the periocular route [31, 32]. Therefore, EVs secreted from retinal organoids could potentially offer a protective cell-free therapy in ocular diseases including the most commonly prevalent AMD, RP, glaucoma, diabetic retinopathy, etc. [33, 34]. Alternatively, retinal organoid-derived EVs could provide insight into disease pathology by serving as biomarkers to monitor the disease onset and/or progression [35].

Atomic force microscope (AFM) has become an attractive characterization tool as it is a label free, non-invasive technique ideal for soft biological samples such as cells, tissues including the EVs [36–38]. The state-of-the-art technology allows for both morphological and nanomechanical characterizations of these samples in fluid environment, thus preserving their overall characteristics [39]. AFM has been used to evaluate the morphology of mesenchymal stem cell (MSC)-derived EVs ornated with LJM-3064 aptamer, a myelin-specific DNA aptamer on remyelination processes and immunomodulatory activity in myelin oligodendrocyte glycoprotein (MOG35-55)-induced mouse multiple sclerosis model [39]. AFM was employed to evaluate the therapeutic platform in MSC derived EVs loaded with doxorubicin against the colorectal cancer, in which, it was again used to characterize their morphology [40]. However, in both these studies, the AFM characterization of EVs was performed in air medium by drying the sample which often results in unrealistic characteristic attributes of biological samples.

In this study, we report the efficient isolation of EVs from the developing 3D retinal organoids differentiated from hiPSC and cultured in a novel PBS Vertical Wheel (PBS-VW) bioreactor. We subsequently characterized and compared the biophysical, nanostructural, nanomechanical, molecular, and proteomic profiling of EVs derived from hiPSC-differentiated retinal organoids with the human umbilical cord mesenchymal stem cells (hUCMSC)-derived EVs, using AFM and proteomics tools. Such wide-range of characterization of retinal organoid EVs are vital to allow their utilization for diagnostic and therapeutic studies.

## 2. Materials and Methods

### 2.1 Human iPSC culturing and retinal organoid differentiation

GFP-knocked in hiPSC [41] and an in-house reprogrammed hiPSC line from peripheral blood using CytoTune™-iPS 2.0 Sendai Reprogramming Kit (A16517, ThermoFisher Scientific, USA) were utilized for this study. hiPSCs were cultured in Matrigel-coated 6-well plate (354234, Corning, USA) with mTeSR™ Plus medium (100-0274, Stemcell Technologies, Canada) until sub-confluency. The cells were washed and lifted using gentle cell dissociation reagent comprising of DPBS (21-031-CV, Corning, USA) containing 0.5% 0.5M EDTA (46-034-Cl, Corning, USA). The lifted small clumps of hiPSC aggregates were cultured in 6-well suspension plate for the generation of retinal organoids *via* the embryoid body (EB) approach as schematically outlined in **Figure 1A** and as per our previously published study [42].

**Figure 1.**
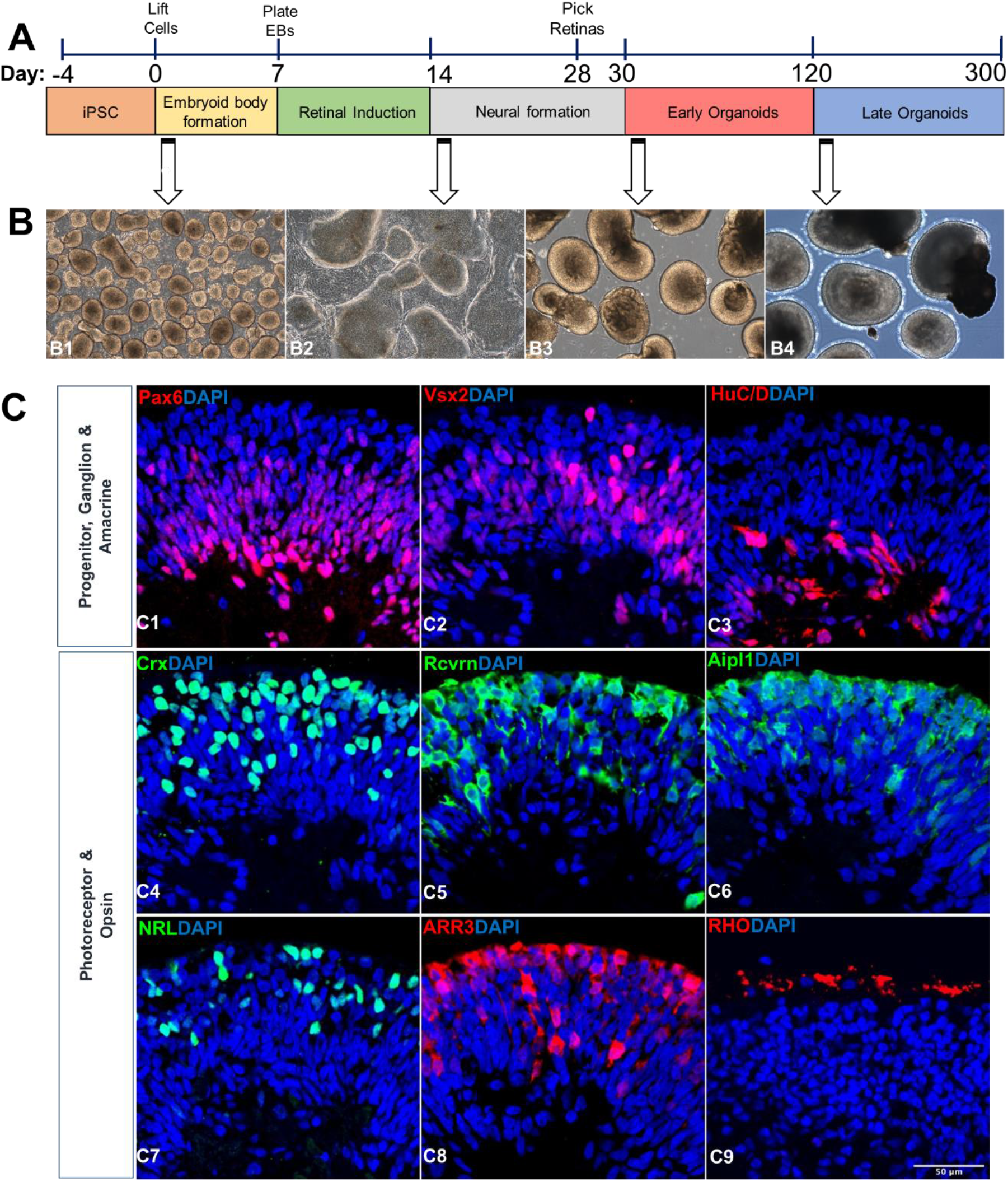
Retinal organoids differentiation and characterization. **(A)** Schematic illustration showing the differentiation of retinal organoids from hiPSC and timeline representation of the early (Day 30-120) and late organoids (Day 121-300) for exosome preparation. **(B)** Phase contrast microscopic images showing the morphology of retinal organoids at different time points of differentiation (B1, Day 2; B2, Day 25; B3, Day 45; B4, Day 200). Scale bar, 200 µm. **(C)** Confocal images of 90-day old retinal organoids (C1-8) stained for retinal progenitor cells (Pax6, C1; Vx2, C2), ganglion and amacrine cells (Brn3a, C3), pan photoreceptors (Crx, C4; Rcvrn, C5; Aipl1; C6), rod photoreceptor (NRL; C7), and cone photoreceptor (ARR3, C8). Image of 200-day old retinal organoids showing the staining of rhodopsin protein indicating the maturation of rod photoreceptor (Rho, C9). Scale bar, 50 µm.

The first day of differentiation is annotated as day 0, with 3:1 of mTeSR™ Plus to neural induction medium (NIM), consisting of DMEM/F12 (SH30023.02, Hyclone, USA), 1X antibiotic/antimycotic solution (SV30079.01, Hyclone, USA), 1X minimum non-essential amino acids (25-025-Cl, Corning, USA), 1X GlutaMAX™ (35050-061, Gibco, USA), 2 µg/mL heparin sulfate (H3149, Sigma, USA), and 1X N2 supplement (17502-001, Gibco, USA). Medium change was done with mTeSR™ Plus: NIM on day 1 (1:1), day 2 (1:3) and day 3-5 (Full NIM). At Day 7, Organized EBs with well-defined outer rim, visible to eyes were further plated onto adhesive (2D) environment for the formation of neuroepithelium using Matrigel-coated plate. Cultures were briefly treated with varying concentrations of human BMP-4 (120-05ET, Peprotech, USA) from day 7 (1.5 nM), day 9 (0.75 nM), and day 12 (0.375 nM) in NIM respectively for the development of neural retina centers. Cultures were then weaned into retinal differentiation media (RDM) comprising of 1:1 DMEM-High glucose (SH30022.02, Hyclone, USA): DMEM/F12, 1X antibiotic/antimycotic solution, 1X minimum non-essential amino acids, 1X GlutaMAX™, and 1X B27 supplement (17504-001, Gibco, USA) from day 15-30 and fed every 2-3 days. Areas of neural centers with clear optic vesicles were dislodged manually using a sterile microtip between day 28-30 and cultured onto 6-well suspension plate with 3D retinal differentiation medium (3D-RDM) consisting of RDM components along with 5% fetal bovine serum (FBS, S11150, R&D Systems, USA), 0.5X CD Lipid Concentrate (11905-031, Gibco, USA), 200 µM taurine (T0625, Sigma, USA). Medium change was done twice weekly from day 30-300. 1 µM of All trans retinoic acid (R2625, Sigma, USA) was added to the 3D-RDM from day 30-120 and were removed further on (from >day 120) from the medium for photoreceptor maturation.

### 2.2 hUCMSC culturing

Dr. David Meckes, FSU - College of Medicine, Florida State University, provided hUCMSCs of passages 0-2. The growth of hUCMSCs for EV isolation was optimized in PBS-VW bioreactors as shown in our previous studies [28]. hUCMSCs were cultivated in culture media free of EVs, consisting of alpha-MEM (12571063, Life Technologies, USA) with 10% EV-free FBS (S11150, R&D Systems, USA),1% sodium bicarbonate (25080094, ThermoFischer Scientific)), and 1% Penicillin/Streptomycin (15140122, ThermoFischer Scientific, USA). The cells were detached using Trypsin-EDTA (T3924, Sigma Aldrich, USA) at ∼80-90% confluency for bioreactor experiments.

### 2.3 Culturing retinal organoids and hUCMSCs in PBS-VW bioreactors

Approximately, 80 retinal organoids were cultured in a 0.1L PBS-VW bioreactor (PBS Biotech Inc., Camarillo, CA) along with 0.25 g of cytodex-1 microcarriers (C0646,Sigma Aldrich,USA) in EV-free 3D-RDM. hUCMSCs and retinal organoids were cultured in separate bioreactors, each with 60 mL of media and agitated at 25 rpm for 5 minutes with 15 minutes stationary phase for 12 cycles at every 4 hours.

### 2.4 Retinal organoid conditioned medium collection for EV isolation

For EV preparation, we categorized retinal organoids into two phases - Early (50-120-days-old) and late (>120-days-old) organoids based on the development and maturation of photoreceptors. Conditioned media from early and late organoids were collected during every media change (twice weekly) and stored at 4°C. Retinal organoids conditioned media were then treated with polyethylene glycol 6000 (80503, VWR International, USA) to extract the EVs which were further subjected to differential ultracentrifugation processes as described previously [28, 43, 44]. Briefly, the EV-containing conditioned media were centrifuged at 500 xg for 5 min at 4°C. The supernatants were collected into fresh tubes and centrifuged again at 2000 xg for 10 min. The supernatants were then centrifuged at 10,000 xg for 30 min and incubated with 16% PEG 6000 at 1:1, overnight in 4°C on a shaker. The media were centrifuged at 3000 xg for 70 min. The supernatants were discarded, and the pellets were resuspended with phosphate buffer saline (PBS) MT21040CV, Fischer Scientific, USA) and ultracentrifuged at 100,000 xg for 70 min. The supernatants were discarded, and the pellets were resuspended in sterile PBS for further characterizations.

### 2.5 Immunohistochemistry (IHC)

Organoids (90 and 200-days-old) were fixed in 4% paraformaldehyde (157-8, Electron Microscopy Sciences, USA) at 4°C for 20 min. After washing with 1X DPBS (3X, 5 min), organoids were equilibrated sequentially in 15% and 30% sucrose for 1 hour respectively until they sink to the bottom. Organoids were embedded in 2:1 of 20% sucrose and OCT Tissue Tek (Sakura Finetek USA, Inc) and flash frozen. Further, 10 µm tissue sections were collected onto Superfrost Plus Microscopic slides (48311-703, Fisher Scientific, USA) and stored at −20°C until use. Sections were pap-pen and permeabilized with 0.1% Triton X-100 (T8787, Sigma Aldrich, USA) in 10% normal donkey serum (NDS) (S30-100ML, Millipore, USA) made in PBS for 15 min. Blocking was done for 1 hour with 10% NDS and sections were incubated with primary antibodies diluted in 10% NDS overnight at 4°C. Complete list of primary antibodies, sources and concentrations are listed in **Suppl. Table S1A**. Following the overnight incubation on the next day, sections were washed with DPBS (3 times with 5 min interval). Species-specific fluorophore-conjugated secondary antibodies diluted at 1:250 in 10% NDS were added onto the sections and incubated for 1 hour in the dark at room temperature. List of secondary antibodies and their sources are listed in **Suppl. Table S1B**. Sections were counterstained for nuclei with 1ug/ml DAPI (10236276001, Roche USA) for 10 min in the dark at the room temperature. The slides were washed thrice with PBS (3 times with 5 min interval), and mounted using Fluoromount-G (17984-25, Electron Microscopy Sciences, USA). High resolution images were acquired with an LSM 700 Laser Scanning Confocal Microscope (Carl Zeiss, USA) and Zen software. Images were processed using Image J (NIH, USA) and adjusted for brightness and contrast.

### 2.6 Particle size, zeta potential and protein assay

Particle size and zeta potential of EVs were assessed using Nanoparticle Tracking analysis (NTA) and Zeta View instrument (ZetaView® TWIN PMX-220) respectively, which utilizes dynamic light scattering (DLS) technique at 25°C with 90° scattering angle. Zeta-View Analysis software was used for data processing as previously explained [all the EV samples were evaluated in triplicates and prepared by diluting with PBS (particle-free)] at 1:1000 [45, 46]. EV protein content was estimated using Pierce™ BCA Protein Assay Kit (23221, ThermoFischer Scientific USA).

### 2.7 Morphological examination by Atomic Force Microscopy (AFM)

Freshly cleaved mica surface was employed to study the morphology and nanomechanical attributes of exosomes under various treatments. Foremost, for effective adsorption of exosomes on mica surface, freshly cleaved mica was modified using a 3:1 mixture (by volume) comprising of 3-aminopropyltriethoxysilane (APTES) and N, N-diisopropylethylamine (DIPEA) for 2 hours at 60°C and used immediately [47]. Working EV sample was prepared by diluting the original EV stock solution 1000-fold with Milli Q water solution. 5 µL of working solution was drop-casted on the APTES modified mica. After 30 min, the mica surface was rinsed with Milli Q water to remove any unbound exosomes. AFM tip calibrations and experiments were then performed in fluid environments using a Dimension Icon Scanasyst AFM (Bruker Corporation, Santa Barbara, CA). Peak Force Quantitative Nanomechanical Mapping (PF-QNM) and nanoindentation experimental technique were employed to study morphology and nanomechanical attributes of the EVs, respectively. A Scanasyst-Air probe with pyramidal tip geometry was used to achieve high resolution topography of the EVs. This probe bearing a sharp tip has a radius of 5 nm and a stiffer cantilever with nominal spring constant of 0.4 N/m. For characterization of topographical morphology, the probe was not calibrated. Topography was assessed using a low peak force of 300 pN and a scan rate of 0.1 Hz. Nanoscope analysis v1.9 software was employed to analyze the morphology characteristics such as height and surface roughness of the EVs. The surface roughness was derived as the root mean square variation in the sample topography. For nanomechanical properties characterization, AFM probe was calibrated in fluid environment on a plain mica surface to determine the spring constant of 0.38 N/m and deflection sensitivity of 35 nm/V. Nanoindentation experiment was performed on at least 50 samples in which, each tip-sample interaction resulted into a force-separation (F-S) curve. A trigger force of 700 pN was applied to engage the tip with the sample. Further, by employing DMT model, each F-S curve was analyzed to yield various nanomechanical attributes such as Young’s modulus (YM), deformation and adhesion.

### 2.8 Quantitative real-time polymerase chain reaction (qRT-PCR)

Total RNA was extracted from 90 and 200-days-old retinal organoids using RNeasy® Mini Kit (74104, Qiagen, USA) as per the instructions of the manufacturer. The cDNA was synthesized using iScript cDNA Synthesis Kit (1708891, Bio-Rad, USA). Real-time qPCR was performed on CFX96 system (Bio-Rad, USA) using iTaq™ Universal SYBR® Green Supermix (64047467, Bio-Rad, USA) to measure the expression levels of EV markers using the primer sequences listed in **Supple. Table S2**. The amplification reactions were performed as follows: 95°C for 30 s, 40 cycles at 95°C for 5 s, 60°C for 25 s, 95°C for 5 s and final extension at 95°C for 5 s. The Ct values of the target genes were first normalized to the endogenous control beta actin. The corrected Ct values were then utilized to compare and validate the exosome biogenesis in late versus early organoids. Log_2_ fold change in gene expression of all targets were then calculated.

### 2.9 Western Blotting of EVs

EV pellets resuspended in 1X PBS, were lysed in equal volume of urea-based lysis buffer. Samples were heated for 5 min at 95°C, and equal amounts (10 µg) were resolved on a 10% running/4% stacking SDS-PAGE gel for 45 min at 100 V until the loading dye entered the resolving layer, then increased to 150 V until the run was completed. The proteins were transferred onto PVDF/ nitrocellulose membrane (1620174, Transbloto, BioRad, USA) and the blots were blocked with 5% BSA (A5611, Sigma Aldrich, USA) for 1 hour. The membranes were incubated overnight at 4°C with primary antibodies - Flotillin 2, CD63, Alix, HRS, Calnexin, Caveolin, and HSP 70 (**Table 1A**) at 1:1000 dilution in 3% BSA solution. Blots were washed with PBST (3X, 5 min) and incubated for 1 hour at room temperature with host-specific secondary antibodies as shown in **Table 1B** (anti-rabbit and anti-mouse) diluted in blocking buffer. The blots were washed with PBST (3X, 5 min) and imaged with a chemiluminescent substrate (AC2101, Azure Biosystems, Dublin, CA, USA) using IMAGEQUANT LAS400 (GE Healthcare Life Sciences, Pittsburgh, PA, USA).

**Table 1.**
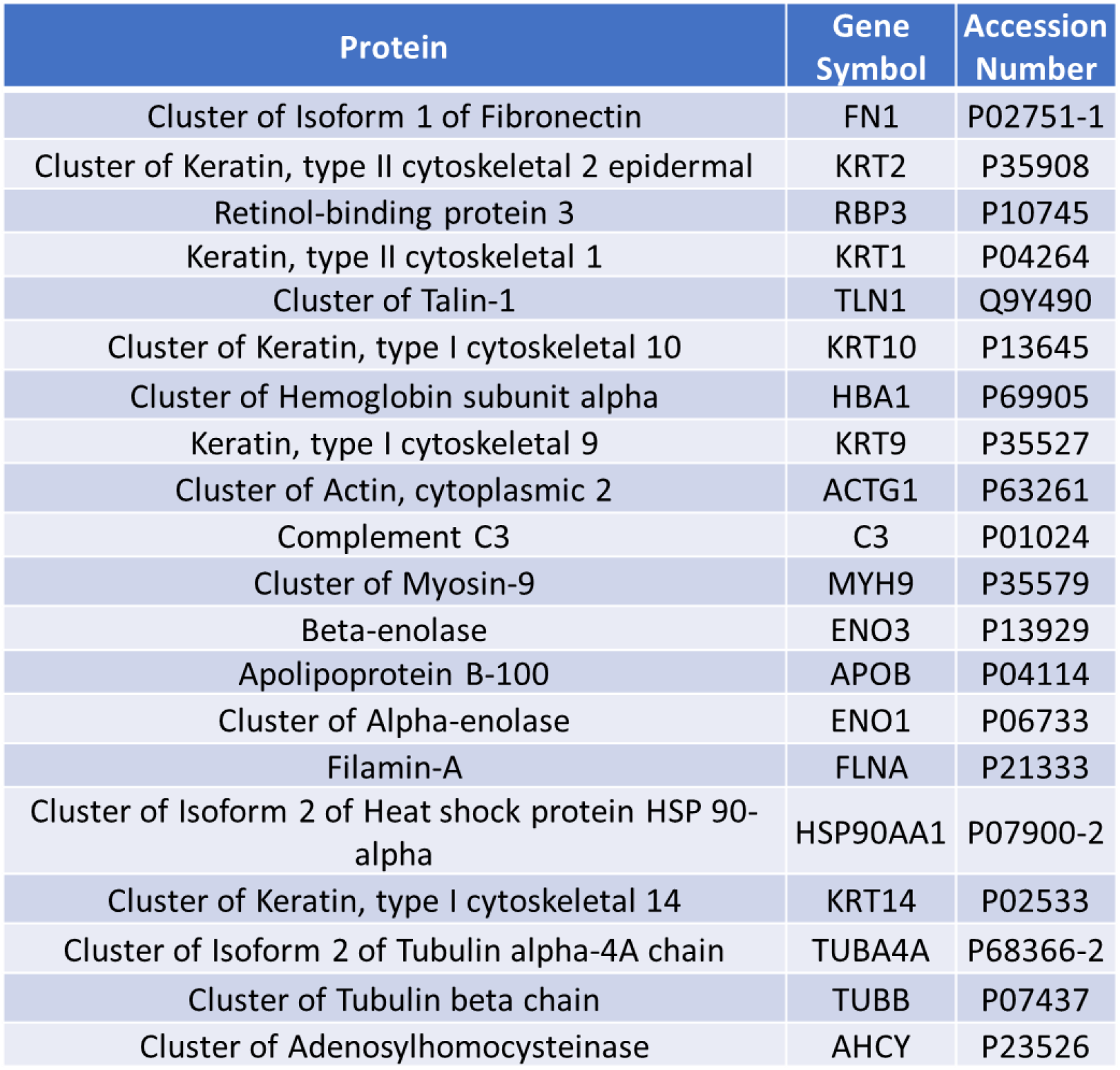
Top 20 identified proteins in the proteomic data.

| Protein | Gene Symbol | Accession Number |
| --- | --- | --- |
| Cluster of Isoform 1 of Fibronectin | FN1 | P02751-1 |
| Cluster of Keratin, type II cytoskeletal 2 epidermal | KRT2 | P35908 |
| Retinol-binding protein 3 | RBP3 | P10745 |
| Keratin, type II cytoskeletal 1 | KRT1 | P04264 |
| Cluster of Talin-1 | TLN1 | Q9Y490 |
| Cluster of Keratin, type I cytoskeletal 10 | KRT10 | P13645 |
| Cluster of Hemoglobin subunit alpha | HBA1 | P69905 |
| Keratin, type I cytoskeletal 9 | KRT9 | P35527 |
| Cluster of Actin, cytoplasmic 2 | ACTG1 | P63261 |
| Complement C3 | C3 | P01024 |
| Cluster of Myosin-9 | MYH9 | P35579 |
| Beta-enolase | ENO3 | P13929 |
| Apolipoprotein B-100 | APOB | P04114 |
| Cluster of Alpha-enolase | ENO1 | P06733 |
| Filamin-A | FLNA | P21333 |
| Cluster of Isoform 2 of Heat shock protein HSP 90-alpha | HSP90AA1 | P07900-2 |
| Cluster of Keratin, type I cytoskeletal 14 | KRT14 | P02533 |
| Cluster of Isoform 2 of Tubulin alpha-4A chain | TUBA4A | P68366-2 |
| Cluster of Tubulin beta chain | TUBB | P07437 |
| Cluster of Adenosylhomocysteinase | AHCY | P23526 |

### 2.10 Mass spectrometry and Proteomics data analysis

Proteins from PB VW bioreactor cultured retinal organoid EVs were purified and digested on S-trap micro column following manufacture’s instruction (C02-micro-10, Protifi, USA). Briefly, 25 µg of EV samples were lysed with 2X SDS lysis buffer (10% SDS, 100 mM TEAB pH 8.5), reduced with 20 mM dithiothreitol (V3155, Promega, USA) at 95℃ for 10 min and alkylated with 40 mM iodoacetamide (RPN6302V, Amresco, USA) at room temperature in dark for 30 min. After acidification, samples were bound in S-trap micro column and washed three times with binding/wash buffer. Digestion buffer with 2.5 µg of sequencing grade Trypsin and ProteinaseMax (V5111, V2071, Promega, USA) was loaded on the column and incubated at 37℃ overnight. On the following day, digested peptides were sequentially eluted with 50 mM Tetraethylammonium bromide (TEAB), 0.2% formic acid, 0.2% formic acid in 50% acetonitrile (MeCN) (89871C, ThermoFisher, USA) and pooled.

The eluted peptides from either methods were submitted to FSU College of medicine Translational Science Laboratory to be analyzed on the Thermo Q Exactive HF (High-resolution electrospray tandem mass spectrometer). Resulting raw files were analyzed with Proteome Discoverer 2.4 using SequestHT, Mascot and Amanda as search engines. Scaffold (version 4.10) software was used to validate the protein and peptide identity. Peptide identity was accepted if Scaffold Local FDR algorithm demonstrated a probability of >95.0%. Likewise, protein identity was accepted if the probability level was >99.0% and contained a minimum of two recognized peptides as previously described [48].

### 2.11. Statistical Analysis

The data values were provided as mean ± SEM. The intergroup differences for all the analysis (except qPCR) were determined with the Graph Pad Prism, version 5.01, using a two-tailed student’s t-test or one-way ANOVA followed by “Bonferroni’s Multiple Comparison Test.” The results were considered statistically significant if the p value was less than 0.05. For gene expression analysis by qPCR, graphs were plotted using Graph Pad Prism, Version 9 using two-way ANOVA.

## 3. Results

### 3.1 Human 3D retinal organoids express retinal markers and opsins

3D retinal organoid cultures recapitulate the developmental timeline of human retinogenesis by encompassing all the various retinal cell types present *in vivo*. Retinal organoids were differentiated using our previously published protocols [42] (**Figure 1A)**. Retinal organoids were easily identified by phase contrast live imaging (**Figure 1B**) displaying typical features of neuroepithelium formation (1B1-B2) and visible laminated pattern (1B3). Sequential development of all retinal cell types took place in the precise order followed by the maturation of photoreceptors exhibiting a brush-like projections (a.k.a. outer segments) on the periphery of the retinal organoids (**Figure 1B4**). Following aggregation as organized EBs within the 1^st^ week of differentiation (**Figure 1B1**) and replating on the Matrigel-coated surface, optic vesicle-like structures were observed in the 2-3^rd^ week of differentiation. The borders of these optic vesicles have shiny edges (**Figure 1B2**), clearly discriminating from the surrounding retinal pigment epithelium and other cells. These were manually lifted 4-weeks following the start of differentiation and cultured as retinal organoids in 6-well suspension plates. Histological examination of retinal organoids by IHC following 90 days of culture identified proteins specific to retinal cell types including neural progenitor cells (PAX6, VSX2; **Figure 1C1-2**), ganglion and amacrine cells (HUC/D; **Figure 1C3**), photoreceptors (CRX, RCVRN, AIPL1, NRL; **Figure 1C4-7**) along with phototransduction protein in cone photoreceptor (ARR3; **Figure 1C8**). The expression of Rhodopsin, a key light sensing protein in rod photoreceptor was detected in the 200-days old mature retinal organoids (RHO, **Figure 1C9**).

### 3.2 Bioreactor-grown retinal organoids releases EVs with superior biophysical properties

EVs were released in the culture medium of both plate and bioreactor-grown (**Figure 2A**) retinal organoids and hUCMSC. The NTA showed that the EV mean diameter was comparable in the late retinal organoid EVs (plate), bioreactor retinal organoid EVs (late), hUCMSC EVs (flask) and hUCMSC EVs (bioreactor) (100.9 ±1.1 nm, 109.1 ± 1.2 nm, 94.2 ±1.7 nm and 97.8 ± 2.3 nm respectively) (**Figure 2C**). The zeta potential for the retinal organoid EVs (plate and bioreactor) and hUCMSC EVs (flask and bioreactor) were −8.47± 0.09mV, −17.32 ±0.2mV, _-_9.30 ± 0.56 mV, and _-_10.49 ± 0.17 mV respectively. The EV particle number per mL of conditioned medium in bioreactor-grown retinal organoid EVs (2.1x 10^11^) was significantly higher (P<0.005) as compared to EVs cultured in plate (9.2×10^10^) (**Figure 2B and 2D)**. In parallel, the increased EV concentration was noticeable in bioreactor-grown hUCMSC with respect to the flask-grown (P<0.005) (**Figure 2B and 2D)**. Similarly, the total protein content estimated by BCA assay showed a considerable increase (P<0.005) in the protein content per mL of conditioned medium in bioreactor-grown retinal organoids EVs (6.26 µg/mL) compared to plate-grown EVs (2.27 µg/mL) (**Figure 2E)**.

**Figure 2.**
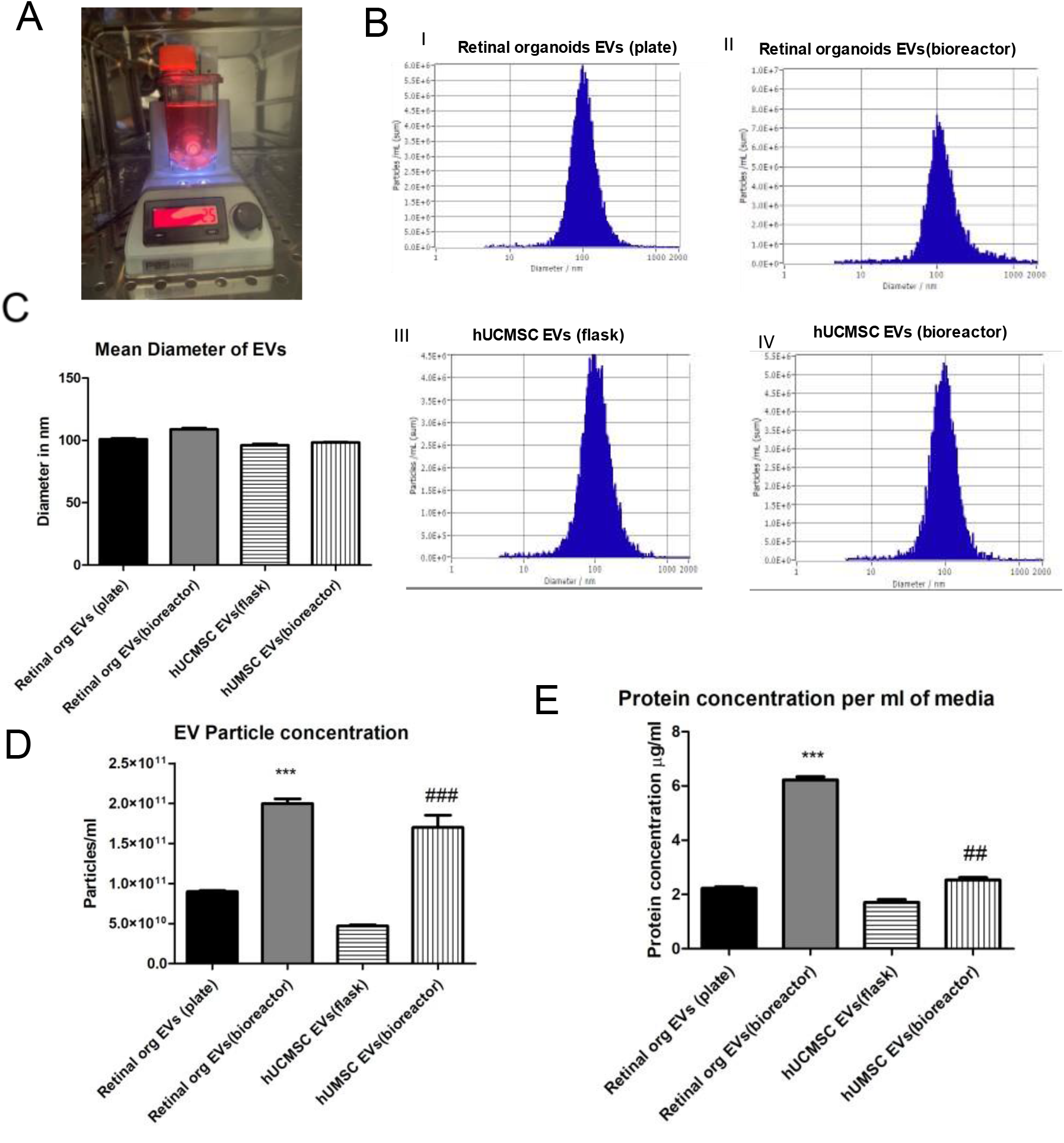
Characterization of Retinal organoid derived EVs by NTA. **(A)** Retinal organoids in PBS VW bioreactor. **(B)** Comparison of EVs from retinal organoids (plate and bioreactor) and hUCMSC (flask and bioreactor) culture. Representative NTA histogram for i) retinal organoids EVs (plate) ii) retinal organoids EVs (bioreactor) iii) hUCMSC EVs (flask) iv) hUCMSC EVs (bioreactor). **(C)** Mean diameter of EVs in nm. **(D)** Retinal organoid and hUCMSCs EV particle concentration per mL of PBS **(E)** Protein content per mL of conditioned medium (* retinal organoid EVs(plate vs bioreactor), # hUCMSC EVs(flask vs bioreactor), *,# indicate p<0.05, **, ## indicate p<0.01 and ***, ### p<0.001

### 3.3 Retinal organoid EVs displayed larger height at Nano structural and softer nanomechanical characteristics

AFM was employed to evaluate the topography of freshly cleaved mica surface as well as post APTES: DIPEA treatment [49]. Both height and peak force error image of freshly cleaved mica surface and APTES: DIPEA treated surface showed no significant change in the topography (**Figures 3, A1-2, and A3-4**). Representative image showing the height and peak force error of EVs derived from retinal organoids (plate and bioreactor), and hUCMSC (flask and bioreactor) are shown at the nanostructural levels respectively (**Figure 3A, 5-6 and 7-8)**. Height images from both hUCMSC and retinal organoids EVs were further quantified to evaluate the height profile and average surface roughness. Retinal organoids EVs displayed significantly larger height (87.95±11.82 nm) than hUCMSC EVs (42.07±5.83 nm) (**Figure 3Ai**). Furthermore, retinal organoid EVs bore significantly more topographical variations (7.39±1.12 nm) in their surface features compared to hUCMSC-derived EVs (2.84±0.33 nm) (**Figure 3 Aii)**. In other words, hUCMSC-derived EVs were smoother compared to retinal organoid-derived EVs.

**Figure 3.**
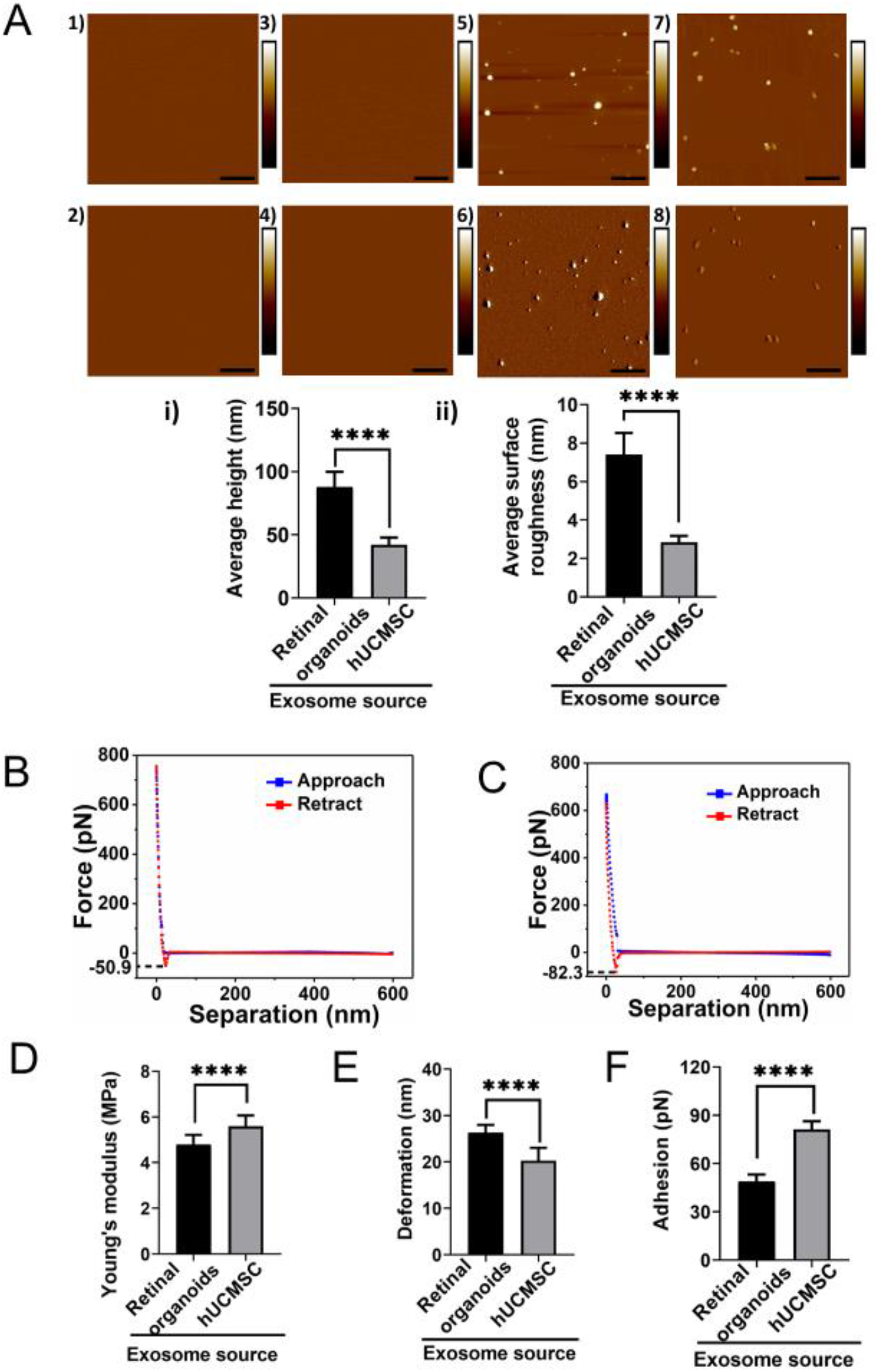
Topographical and nanomechanical characteristics of EVs. **(A)** Representative AFM images of plain mica and APTES: DIPEA modified mica (1, 3) Height image (2) Peak force error image. Representative images of retinal organoid and hUCMSC EVs (5, 7) Height image 6, 8) Peak force error image. (Scale bar: 1µm; color bar for height image: −65 nm to 65 nm; color bar for peak force error image: −300 pN to 300 pN). Morphology quantification i) Average height ii) Average surface roughness. (Statistical significance: ****; p<0.0001). A representative force-separation curve displaying adhesion value corresponding to exosomes derived from **(B)** Retinal organoid and **(C)** hUCMSC. Nanomechanical attributes displaying **(D)** Young’s modulus **(E)** Deformation **(F)** Adhesion. (****; p<0.0001).

Following this, nanoindentation experiments evaluated the nanomechanical attributes of EVs derived from retinal organoids and hUCMSC. In this assessment, each interaction between the AFM tip and the sample surface generated a force-separation (F-S) curve. Representative F-S curves for retinal organoid and hUCMSC-derived EVs are shown in **Figure 3B and C**, respectively. As seen from the figure, blue data line represents the trace motion of the tip where, tip is approaching the sample. Red data line represents the retrace motion of the tip in which, the tip retracts from the sample surface. For analysis purpose, retract curve was evaluated by curve fitting technique with the Derjaguin-Muller-Toporov (DMT) contact mechanics model to quantify sample’s Young’s modulus (Y_M_) along with deformation and adhesion. EVs derived from retinal organoids were significantly softer than the hUCMSCs (**Figure 3D)**. The Y_M_ for EVs derived from retinal organoids and hUCMSC were observed to be 4.79±0.42 MPa and 5.59±0.47 MPa, respectively. On the other hand, for EVs derived from retinal organoids, the deformation was observed to be 26.36±1.63 nm and this was significantly more than hUCMSC-derived EVs which were 20.29±2.74 nm (**Figure 3E)**. Furthermore, the adhesion characteristics of EVs derived from retinal organoids was shown to be significantly lesser (48.94±4.15 pN) compared to hUCMSC-derived EVs (81.27±4.94 pN) (**Figure 3F)**. Overall, these nanomechanical attributes could potentially serve as distinguishing parameters to identify and differentiate the EVs derived from retinal organoids and hUCMSC.

### 3.4 Enhanced EV biogenesis was presented by matured retinal organoids

Biogenesis of EVs are modulated and regulated by the molecular machinery of the cells including genes-related to the endosomal sorting complex required for transport proteins (ESCRT)-dependent (Alix, TSG101), and ESCRT-independent proteins (synthenin1 and syndecan1). Rab family of small GTPase proteins including the Rab27 modulates the transport, fusion, and secretion of EVs. Gene expression analysis of retinal organoids collected at early (Day 90) and late (Day 200) time-point showed significantly higher log_2_ fold change of ESCRT (dependent and independent) and Rab targets including *TSG101, ALIX and RAB* from the older organoids implicating abundant synthesis of EVs (**Figure 4A**). Members of tetraspanin family including CD63, CD81, being the pan targets of EV genes were also ∼1-1.4 log_2_ fold higher in older organoids relative to the younger ones. No significant change was seen in the ESCRT-independent targets including syndecan1 and synthenin1. Expression level of Adam10, another EV marker remained unchanged during retinal differentiation.

**Figure 4.**
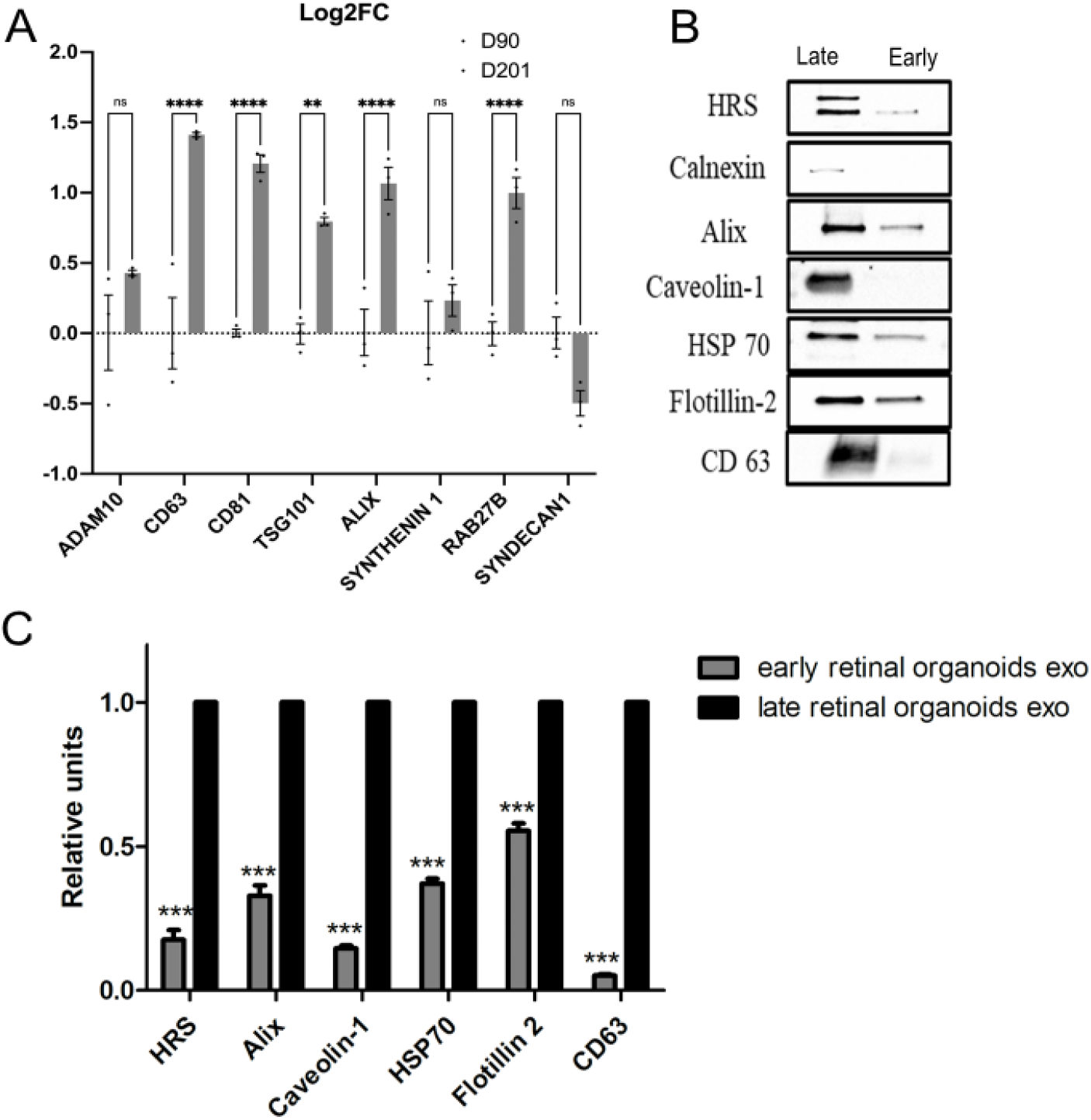
Biogenesis of exosome in human 3D retinal organoids. **(A)** qRT-PCR assay of early (Day 90) and late (Day 200) retinal organoids showing the expression of ESCRT-dependent (Alix, TSG101), ESCRT-independent (Synthenin1, Syndecan1), Rabs family of transport and membrane fusion and pan exosome targets (Rab27B, CD63, CD81, ADAM10). (ns, p>0.05; *p≤0.05; **p≤0.01; ***p≤0.001; ****p≤0.0001). **(B)** Western blots of early and late retinal organoid EVs for exosome markers HRS, Calnexin, Alix, Caveolin 1, HSP 70, Flotillin-2 and CD63. **(C)** Densitometric analysis of western blots. ***represents p<0.001.

### 3.5 Retinal organoid-EVs expressed exosomal marker proteins ubiquitously throughout the development and maturation phases

EVs derived from conditioned medium of young and older retinal organoids have been reported to express ESCRT-dependent proteins, essential for protein trafficking and exosome production [19, 50], and CD63, a tetraspanin surface marker. In this study, EVs derived from older retinal organoids culture showed a higher expression of other markers including HRS (100 kDa), Alix (95 kDa), Caveolin-1 (21kDa), HSP70 (70 kDa), Flotillin-2 (47 kDa) and CD63 (25 kDa) as compared to young retinal organoids-derived EVs, P<0.005 **(Figure 4B and 4C)**. Calnexin (90 kDa), an intracellular protein contamination was completely absent in EVs of younger organoids as opposed to some detection of it from older retinal organoids. This suggests that there has been a ubiquitous release of EVs throughout the development phase of retinal organoids with enhanced expression of EV marker proteins during the maturation phase of older retinal organoids.

### 3.6 Differentially expressed signature proteins were expressed in retinal organoid EVs

To obtain a complete protein profiling of the retinal organoid EV, we preformed proteomics and used hUCMSC EV as control group for comparison. There was a total of 1389 differentially expressed proteins (DEPs) identified, and about 91.8% (1275/1389) DEPs were overlapped (**Figure 5A**). Through Gene Ontology (GO) analysis, 410 DEPs related to retinal homeostasis, neurogenesis, angiogenesis, and interaction with extracellular matrices (ECM) were identified (**Figure 5B**). Proteins from the retinal organoid EV also expressed 16 signature overlaps with retinal tissue enriched proteomic database of Human Protein ATLAS (**Figure 5C**). This includes RBP3 (Retinol-binding protein 3) which is identified in the top 20 of all the proteins (**Table 1**). A manual search of all the protein list was also conducted; we found more retinal function related proteins and EV biogenesis/marker proteins (**Suppl. Table S3, S4**), which provide evidence that EV isolated from retinal organoids are very likely to have superior therapeutic effect on retinal diseases than hUCMSC-derived EVs. Based on the literature [51–53], hMSCs have protective effects on damaged retina regeneration by their unique secretome with immunomodulatory, angiogenesis, anti-apoptotic and neurotrophic factors. We found some growth factors (PDGF), its receptor (PDGFR), and other carrier/transporting proteins (ABCA4, BSG, CDHR1, SLC2A1) were included in the highly expressed list of retinal EVs (**Table 2**). Lower level of Programmed cell death 4 (PDCD4) identified has been beneficial based on the previous findings [51]. Altogether the effects of these proteins are beneficial to maintain the viability of healthy photoreceptors.

**Figure 5:**
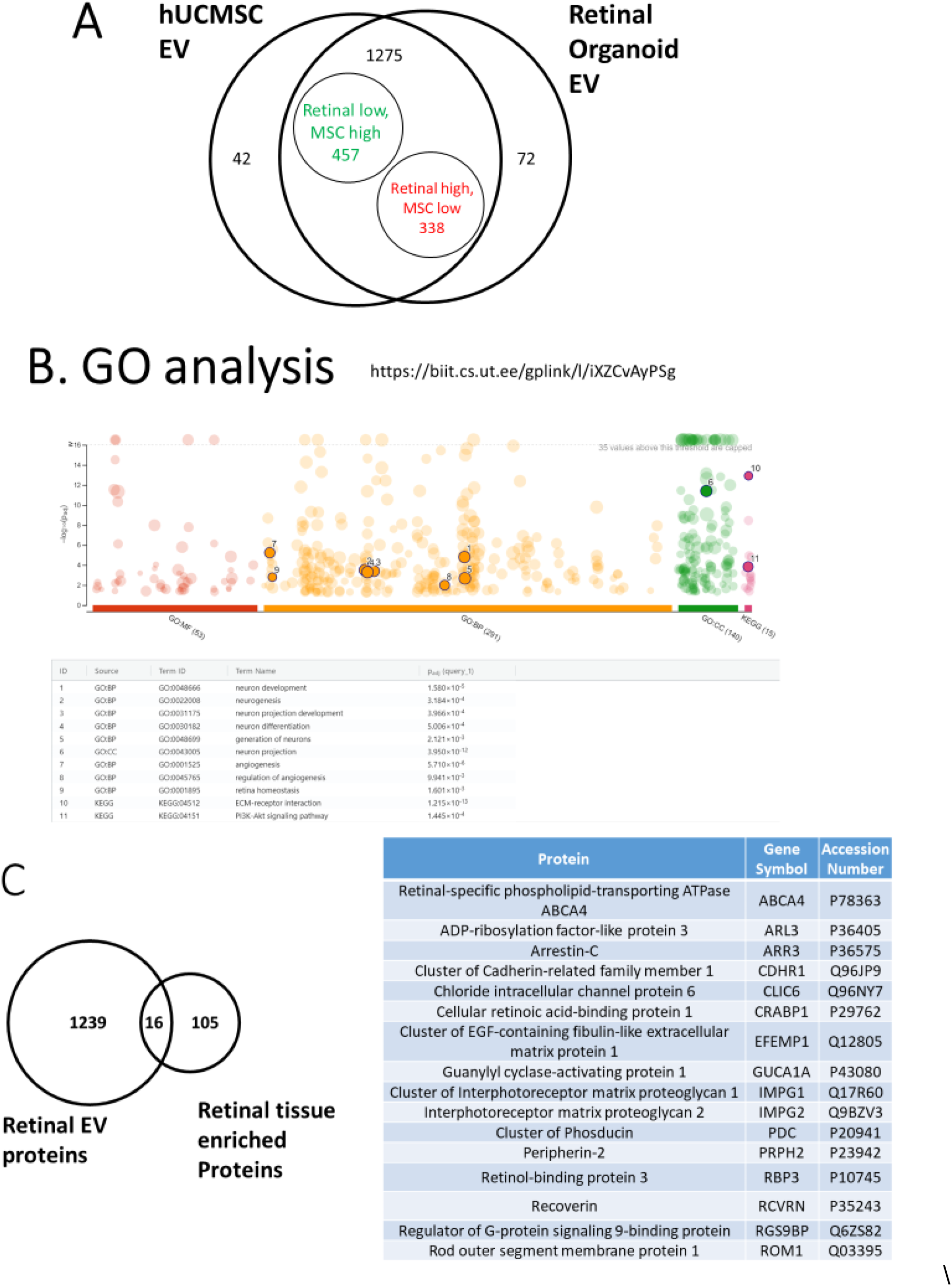
Proteomics analysis comparing retinal organoid with hUCMSC EVs. **(A)** Venn diagram of total identified proteins in both EV groups. **(B)** GO analysis of Retinal EVs. **(C)** Venn diagram of retinal organoid EVs, Human Protein ATLAS of retinal tissue and overlapped proteins between them.

**Table 2.** Proteins related with therapeutic effects.

| Protein | Gene Symbol | T-Test (p-value) | Fold Change |
| --- | --- | --- | --- |
| Cluster of Platelet-derived growth factor C | PDGFC | 0.05 | 2.5 |
| Platelet-derived growth factor receptor beta | PDGFRB | 0.04 | INF |
| Cone cGMP-specific 3',5'-cyclic phosphodiesterase subunit alpha' | PDE6C | 0.07 | INF |
| Isoform 2 of Programmed cell death protein 4 | PDCD4 | 0.01 | 0.0 |
| Cluster of Programmed cell death protein 6 | PDCD6 | 0.23 | 1.4 |
| Retinal-specific phospholipid-transporting ATPase ABCA4 | ABCA4 | < 0.00010 | INF |
| Cluster of Isoform 3 of Basigin | BSG | 0.00 | 2.0 |
| Cluster of Cadherin-related family member 1 | CDHR1 | 0.00 | INF |
| Solute carrier family 2, facilitated glucose transporter member 1 | SLC2A1 | 0.04 | 2.5 |

## 4. Discussion

EVs play a crucial role in intercellular communication by regulating the genetic and proteomic information between the adjacent cells as well as distant organs [54]. Although there has been an increased interest in developing the therapeutic potential of retinal organoid-secreted EVs for ocular diseases, knowledge in this field is still limited. This is one amongst the first study to characterize the protein cargo of retinal organoid-derived EVs using a PBS-vertical wheel bioreactor. A recent study by Zhou et al demonstrated the release of EVs in the conditioned medium and comprehensively characterized the genetic cargo of EVs from three developing time-point of 3D human retinal organoids (D42, D63, and D90) [29]. Uptake of the EVs secreted by the retinal progenitor cells in the early time-point retinal organoids has shown to regulate the gene expression of early differentiated retinal cell types including ganglion cells, amacrine cells, horizontal cells and photoreceptor progenitors and precursors. Other late differentiated retinal cell types beyond the D90 time point includes bipolar and Müller glia. Most importantly maturation of photoreceptor is the final and significant event of human retinogenesis. Most inherited retinal degeneration gets initiated with the degeneration of outer segments followed by the complete loss of photoreceptors. Hence, our study focused on profiling the late time-point retinal organoids, where the photoreceptors are completely mature with inner/outer segments. Additionally, we employed the use of bioreactors to upscale the secretion of EVs by the retinal organoids. The EV profiling of conditioned medium across the complete development and maturation of retinogenesis can serve as a valuable biomarker for assessing the patient-specific retinal organoids with inherited retinal degenerations.

As stated above, EVs likely have a critical role in human retinal development and disease. Klingeborn et al. summarized various studies on the functions of EVs in the healthy and diseased eye, including AMD, diabetic retinopathy, and uveal melanoma [55]. A study showed that MSC-derived EVs administered intravitreally effectively improved hyperoxia-induced retinopathy. Intravitreal administration of microglia-derived EVs in a mouse model of retinopathy of prematurity (ROP) showed a decrease in avascular areas and neovascular tufts, reduced VEGF and transforming growth factor β (TGF-β) expression and photoreceptor apoptosis inhibition [56].

With the advent of technology enabling AFM studies in fluid environment and its ease of operation, AFM technology is widely preferred tool over other techniques such as transmission electron microscope, scanning electron microscope, optical microscope as well as micropipette aspiration [57]. One of the advantages of AFM morphology study is that it yields a 3D topography of the EV surface, which enables its quantification in terms of height profile in addition to surface roughness. Surface roughness indicates the average topographical feature variations on the sample surface and is a measure of its smoothness. Together, height and surface roughness provide complete morphological characterization and serve as quantitative signatures to differentiate between EVs derived from retinal organoids and hUCMSCs. AFM’s height quantification for EVs derived from retinal organoids matches closely with the corresponding NTA data as seen from Figures 2A and 2C. Unlike other conventional techniques, AFM yields various nanomechanical attributes such as Y_M_, deformation and adhesion that can serve as distinguishing signature properties. Y_M_, a measure of sample’s stiffness exhibited that retinal organoids EVs are significantly softer than those derived from hUCMSC and can be attributed to their source. Deformation attribute is often observed to be complimentary to the Y_M_ and is due to the resistance offered by the sample surface to the applied trigger on the AFM tip provided same force is applied. Adhesion is another nanomechanical attribute arising from the repulsive force the tip experiences just before retracting completely from the sample surface and can be seen from the dip in the force values (usually on the negative y-axis scale) shown in Figure 3B and C. These nanomechanical attributes could indicate that stiffer exosomes surface exerts lesser attractive pull on the AFM tip prior to its retraction.

MSC-derived EVs have been shown to have beneficial effects in a rat model of corneal allograft rejection therapy [59]. Therefore, this study compared the protein cargo of retinal organoid-EVs with hUCMSC-EVs by proteomic profiling. The results show that the retinal organoid-EVs have 16 signatures overlapping with retinal tissue enriched proteomic database of Human Protein ATLAS, which include RBP3 and retinal function-related proteins, as well as EV biogenesis markers, neurotrophic and anti-apoptotic factors etc., in comparison to hUCMSC-EVs. The 3D organoid culture may contribute to the protein cargo alteration, as 3D architecture is more efficient in producing EVs and impact EV content including miRNA and proteins [60–62]. In addition, our previous studies have compared the EVs of undifferentiated hiPSCs and isogenic lineage-specific hiPSCs (e.g., ectoderm differentiation), showing that the EV properties are affected during differentiation stage and lineage specification from the hiPSCs [63, 64]. The retinal induction of hiPSCs should promote the EV cargo profile that enhances the retinal function.

Zhou et al. recently published a study which demonstrated that the functional 3D retinal organoids continually generated EVs containing molecular cargo and are linked to post-translational modification and human retinal development regulation [29]. Further it was suggested that EVs may transport their miRNA cargo to human retinal progenitor cells (hRPCs) and the expected targets involved in different stages of retinal development were also studied. According to this study, the internalization of 3D retinal organoid EVs by hRPCs and active transport throughout cytoplasmic compartments resulted in the control of gene expression. While their study characterized the small RNA and miRNA profiles of EVs secreted by retinal organoids, no protein cargo was reported [29]. In addition, the dynamic culture environment in PBS VW bioreactor system [65] used in our study promotes nutrient/oxygen transfer as well as EV biogenesis, in comparison to the static retinal organoid cultures used in the literature. Taken together, the enhanced therapeutic protein cargo profile in the retinal organoid EVs may be attributed to the unique culture system together with the retinal maturation protocol used in our study.

Prospectively, the retinal organoid EVs characterized in our study can be used for drug loading and drug delivery as carriers, as well as for *in vivo* therapeutics for treating ocular diseases. The functional properties of the retinal organoid EVs are yet to be investigated and the potential for scalable production in the large-scale PBS VW bioreactors are yet to be demonstrated. The roles of the specific proteins in the retinal organoid EV cargo may need to be revealed and the approaches to modify the EV cargo (e.g., protein overexpression and the knockout) still needs to be identified. Nonetheless, this study provides advanced knowledge of retinal organoid EVs characteristics for potential ocular disease treatments.

## Conclusion

There has been increased interest in the therapeutic role of EVs in ocular diseases. This study presents a complete morphological, nanomechanical, molecular, and proteomic characterization of retinal organoid EVs, which is necessary for use in diagnostic and therapeutic studies. Our findings show that EV biogenesis markers are highly expressed in late retinal organoids compared to early retinal organoids. This paper specifically explored the protein cargo of late retinal organoids and hUCMSC-derived EVs. The unique culture system used in conjunction with the retinal maturation process suggests that retinal organoid-EVs have an enhanced therapeutic protein cargo profile which can have potential effects in the treatment of ocular diseases.

## Acknowledgement

This research is partly funded by the National Institute on Minority Health and Health Disparities, Grant/Award Number: U54 MD007582, and NSF-CREST Center for Complex Materials Design for Multidimensional Additive Processing (CoManD), Grant/Award Number:1735968. Work was supported, in part, by grants from the National Institutes of Health (R01-EY032197 and U24 EY029891 to D.A. L, R01-NS125016 to Y.L.) P30 Vision Core grant to UCSF Dept of Ophthalmology (EY002162), the Research to Prevent Blindness (unrestricted grant to UCSF Dept of Ophthalmology) and Eagles fifth District Cancer Telethon−Cancer Research Fund and Jay and Deanie Stein Career Development Award for Cancer Research at Mayo Clinic Jacksonville, 2019 Benefactor Funded Champions for Hope Pancreatic Cancer to S.B.

## Declarations

The authors declare that they have no competing interests.

## Data Availability

The datasets generated and analyzed from the current study is available with the corresponding author on reasonable request. The mass spectrometry proteomics data have been deposited to the ProteomeXchange Consortium *via* the PRIDE partner repository with the dataset identifier.

## Notes

### Competing Interest Statement

The authors have declared no competing interest.

