## Supplementary Table 1,2,3,4 for "Biophysical, Molecular and Proteomic profiling of Human Retinal Organoids derived Exosomes"

**Suppl. Table S1A**. Primary Antibody List.

| Protein Name | Host | Dilution | Source | Cat. No. |
|---|---|---|---|---|
| Aipl1 | Rabbit | 1:1000 | Gift from Ramamurthy Lab, WVU | |
| Alix | Rabbit | 1:1000 | Cell Signaling Technologies | 92880S |
| Arr3 | Mouse | 1:100 | Gift from Peter MacLeish Lab, MSM | |
| Calnexin | Rabbit | 1:1000 | Cell Signaling Technologies | 2679S |
| Caveolin-1 | Rabbit | 1:1000 | Cell Signaling Technologies | 3267S |
| CD63 | Rabbit | 1:1000 | Cell Signaling Technologies | 55051S |
| Crx | Rabbit | 1:250 | Abcam | ab140603 |
| Flotillin-2 | Rabbit | 1:1000 | Cell Signaling Technologies | 3436S |
| HRS | Rabbit | 1:1000 | Cell Signaling Technologies | 15087S |
| HSP70 | Rabbit | 1:1000 | Cell Signaling Technologies | 4873S |
| HuC/D | Mouse | 1:100 | Life Technologies (Invitrogen) | A-21271 |
| NRL | Goat | 1:400 | R&D Systems | AF2945 |
| Pax6 | Mouse | 1:100 | DSHB | PAX6-s |
| Rcvrn | Rabbit | 1:500 | EMD Millipore | AB5585 |
| Rho | Mouse | 1:250 | Millipore | MABN15 |
| Vsx2 / Chx10 | Mouse | 1:100 | Santa Cruz Biotechnology, Inc. | sc-365519 |

**Suppl. Table S1B**. Secondary Antibody List.

| Species | Target | Fluorochrome | Source | Cat. No. |
|---|---|---|---|---|
| Donkey | Anti-mouse | Alexa Fluor 555 | Invitrogen | A-31570 |
| Donkey | Anti-mouse | Alexa Fluor 647 | Invitrogen | A-31571 |
| Donkey | Anti-rabbit | Alexa Fluor 555 | Invitrogen | A-31572 |
| Donkey | Anti-rabbit | Alexa Fluor 647 | Invitrogen | A-31573 |
| Donkey | Anti-Goat | Alexa Fluor 555 | Invitrogen | A-21432 |
| Donkey | Anti-Goat | Alexa Fluor 647 | Invitrogen | A-21447 |
| Goat | Anti-Rabbit | - | Cell Signaling Technologies | 7074 |
| Goat | Anti- Mouse | - | Cell Signaling Technologies | 56970 |

**Abbreviations:** Aipl1, Aryl hydrocarbon receptor interacting protein like 1; Arr3, Arrestin3; Crx, Cone-rod homeobox; HuD, Hu-antigen D; NRL, Neural retina leucine zipper; Pax-6, Paired box protein Pax-6; Rcvrn, Recoverin; Rho, Rhodopsin; Vsx2/Chx10, Visual system homeobox 2/Homeobox protein CHX10.

**Suppl. Table S2**. qPCR Primers.

| Gene Symbol | Forward primer (5'→3') | Reverse primer (5'→3') |
|---|---|---|
| *CD63* | ACAACCACACTGCTTCGATCC | GACTCGGTTCTTCGACATGGA |
| *CD81* | ATCCTGTTTGCCTGTGAGGTG | TGCTGTAGGGCCTGGTCATAG |
| *TSG101* | CACCTGGTGGTCCATATCCTG | GATGGTGTCCTCGCTGATTGT |
| *ALIX* | TAAGTGCATCTGAGGGCCAAA | GGGGCCTCCTTTCCTAGTTTC |
| *SYNTHENIN 1* | TTATTAAGCGGATGGCACCAA | TTGCCAAAGAAGGAAACTGGA |
| *RAB27B* | TCCATGAAGCTGCTTGTCTCA | GTTGGGTCTCCACCCAGAAAT |
| *SDC1* | TCGAATCTCTGTGCCTTCGTC | AAACCTTGGCTGAACCTACCG |
| *ADAM10* | TGGCTACTTCAGCTCCCATTC | TTCCTTCCGCTAGACCCTCAG |
| *BETA ACTIN* | GTACTCCGTGTGGATCGGCG | AAGCATTTGCGGTGGACGATGG |

### Suppl Table S3. Retinal related proteins

| Protein | Gene Symbol | Accession Number |
|---|---|---|
| Retinol-binding protein 3 | RBP3 | P10745 |
| Retinal dehydrogenase 1 | ALDH1A1 | P00352 |
| Cluster of Retinal guanylyl cyclase 1 | GUCY2D | Q02846 |
| Transthyretin | TTR | P02766 |
| Retina-specific copper amine oxidase | AOC2 | O75106 |
| Retinoic acid-induced protein 3 | GPRC5A | Q8NFJ5 |
| Cluster of Retinol-binding protein 1 | RBP1 | P09455 |
| Cellular retinoic acid-binding protein 1 | CRABP1 | P29762 |
| Retinal-specific phospholipid-transporting ATPase ABCA4 | ABCA4 | P78363 |
| Cluster of Isoform 3 of Retinal dehydrogenase 2 | ALDH1A2 | O94788-3 |

### Suppl Table S4. EV related proteins

| Protein | Gene Symbol | Accession Number |
|---|---|---|
| CD81 antigen | CD81 | P60033 |
| CD9 antigen | CD9 | P21926 |
| Tumor susceptibility gene 101 protein | TSG101 | Q99816 |
| Ras-related protein Rab-21 | RAB21 | Q9UL25 |
| Isoform Short of Ras-related protein Rab-27A | RAB27A | P51159-2 |
| Ras-related protein Rab-27B | RAB27B | O00194 |
| Syntenin-1 | SDCBP | O00560 |
| Cluster of Programmed cell death 6-interacting protein | PDCD6IP | Q8WUM4 |
| Cluster of Isoform 2 of Heat shock protein HSP 90-alpha | HSP90AA1 | P07900-2 |
| Flotillin-1 | FLOT1 | O75955 |
| Cluster of Vimentin | VIM | P08670 |

**Supplementary Data file:**

Excel Spreadsheet “Samples view report (hUMSC-CtlR)
